# Evolutionarily divergent DUF4465 domains have a common vitamin B_12_-binding function

**DOI:** 10.1101/2025.08.21.671476

**Authors:** Charlea Clarke, Michal Banasik, Rokas Juodeikis, Martin J. Warren, Richard Pickersgill

## Abstract

Domains of Unknown Function (DUFs) comprise a large portion of the bacterial proteome, yet their biological roles remain poorly understood. We recently identified two DUF4465 proteins (IPR027828 family proteins), BtuJ1 and BtuJ2, in the vitamin B_12_–auxotrophic gut commensal *Bacteroides thetaiotaomicron*, which act as high-affinity B_12_–binding proteins that scavenge the cofactor to ensure survival. Such B_12_ capture is essential for bacteria that have lost the ability to synthesize B_12_ *de novo*. The DUF4465 family contains more than 1,000 members distributed across eight bacterial clades in gut microbiome and marine environments, raising the question of whether B_12_-binding is ubiquitous across this family. Here, we show that B_12_-binding is conserved across five additional sequence-diverse DUF4465 proteins bringing the total we have characterized to seven. Structural and biochemical analyses, including the crystal structure of D5EK51 from *Coraliomargarita akajimensis* bound to B_12_, reveal a conserved augmented β-jellyroll fold and a shared B_12_-binding motif. Together, these findings establish DUF4465 as a structurally conserved family of B_12_-binding proteins and point to their widespread role in microbial competition for this essential cofactor.

## Introduction

Nutrient acquisition is a critical aspect of bacterial survival, particularly for essential micronutrients, such as vitamin B_12_ (cobalamin or B_12_) which are often limited in the environment^1^. This scarcity is especially evident in the human gut, where only a minority of the bacterial species present can synthesize B_12_ completely^2,3^, and where over 80% of sequenced human microbial gut species nevertheless encode B_12_ dependent genes or riboswitches^4,5^. In such ecosystems, microbial communities rely heavily on both sharing and competition for B_12_^6,7^. The importance of this cofactor stems from its central role in metabolism, acting as a cofactor for key enzymes such as methionine synthase and methyl malonyl CoA mutase^8–10^. Cobalamin biosynthesis is among the most complex and energetically demanding metabolic processes known, requiring approximately 30-enzymatic step biosynthetic pathway^11,12^. Although B_12_-independent enzymes and pathways can substitute for survival, most bacteria that encode them also retain B_12_-dependent isoenzymes and rely on transporters to acquire complete or incomplete corrinoids through salvage pathways^13,14^. This pattern suggests that scavenging for B_12_ is not only sufficient but often preferred, as reflected in microbial communities where B_12_ producers are outnumbered by B_12_-dependent organisms such as the gut commensal *Bacteroides thetaiotaomicron*^15–17^.

Recent studies have expanded our understanding of B_12_ transport and sequestration, particularly in B_12_-limited environments such as the human gut. While the core B_12_ uptake pathway is highly conserved, scarcity exerts a strong selective pressure to diversify this protein network. In *Bacteroides thetaiotaomicron*, B_12_ limitation induces expression of additional outer membrane lipoproteins including those encoded by *btuG, btuH*, and *btuJ1/J2*, which are co-located within riboswitch-regulated loci with *btuB, btuCD*, and *btuF* ^18–20^. BtuG, H, J1 and J2 directly bind B_12_ and enhance uptake efficiency, conferring a competitive edge under nutrient limitation^21–23^. Such adaptations highlight how otherwise conserved B_12_ transport systems evolve diversification in response to selective pressure. We recently demonstrated that the DUF4465 proteins BtuJ1 and BtuJ2 are high-affinity B_12_-binding proteins that localize to bacterial extracellular vesicles and promote the survival of *Bacteroides thetaiotaomicron*^24^. Independent work has also reported B_12_-binding to BtuJ1 aids bacterial survival^25^. Here, we show that B_12_ binding is a unifying feature of DUF4465 proteins, demonstrated across seven family members with as little as 16% sequence identity.

## Results

### Comparative Analysis of the DUF4465 family

A phylogenetic analysis of the DUF4465 family revealed a broad distribution across eight bacterial clades (Figure 1A). Notably, sequence variation within the *Bacteroidetes* exceeds that observed between different clades. To capture this diversity, we selected representative proteins of the family, including “D5EK51” from *Coraliomargarita akajimensis*, “Pan” from *Mucisphaera calidilacus* (A0A518BTS0), “Fluta” from *Fluviicola taffensis* (F2I9N3) and “656” from *Rikenellaceae bacterium* (A0A928FT42), “Cylst” from *Cylindrospermum stagnale* PCC 7417 (K9X579), to span the sequence space of this family (Figure 1B-E). Consistent with a role at the cell surface, all representative proteins possess N-terminal signal peptides as predicted by SignalP^26^.

**Figure 1.**
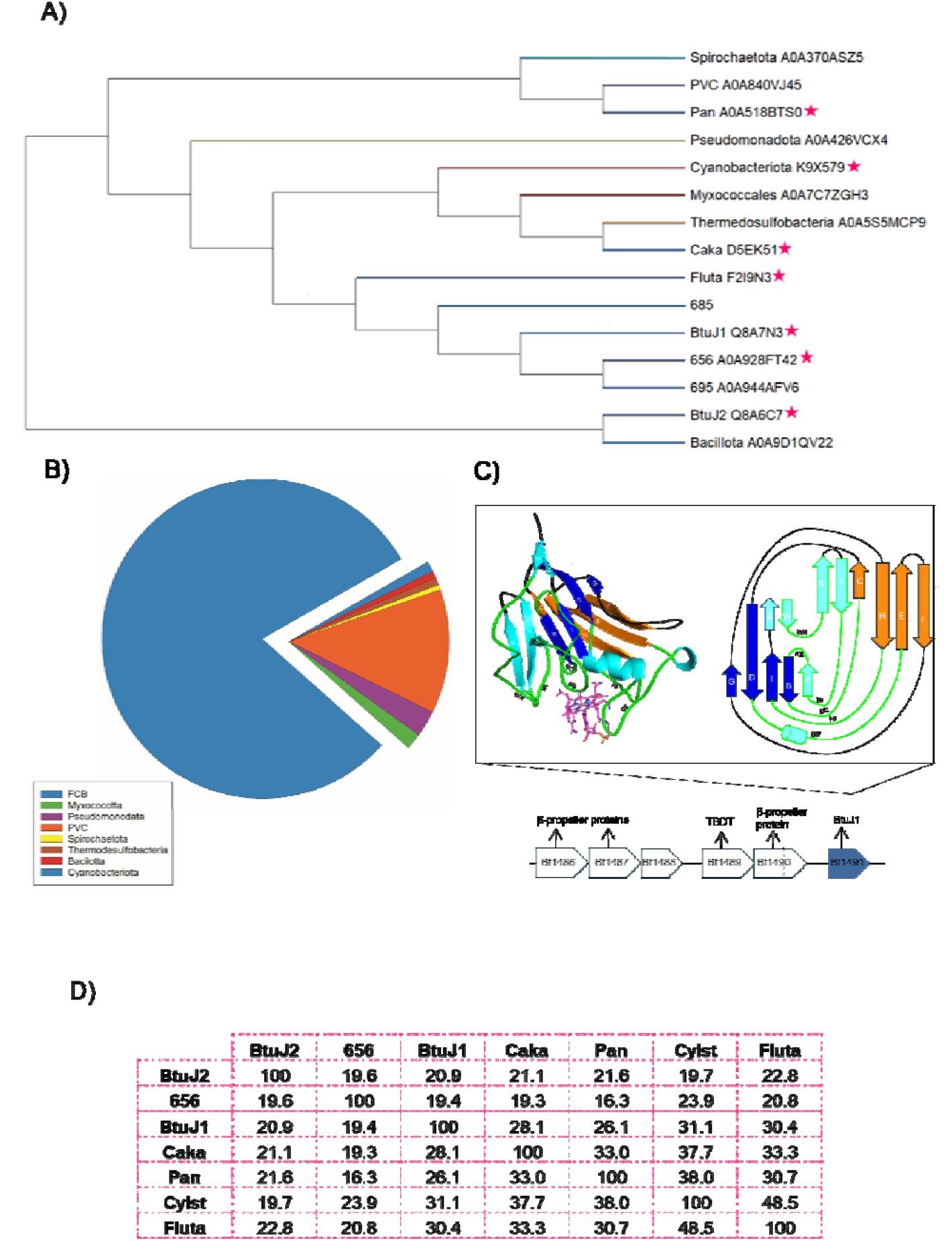
Evolutionary distribution, structural features, and genomic context of DUF4465 family proteins. A) Phylogenetic tree of representative DUF4465 (IPR027828 family) proteins sequences from diverse bacterial phyla. Branch lengths are proportional to sequence divergence. Pink stars indicate proteins selected for study. B) Taxonomic distribution of DUF4465 homologues shown as a proportional pie chart; the FCB group represents the majority of sequences, with smaller contributions from *Myxococcota, Pseudomonadota*, PVC, *Spirochaetota, Thermodesulfobacteria, Bacillota*, and *Cyanobacteriota*. C) To the left is shown the structure of *Bacteroides thetaiotaomicron* DUF4465 protein (gene identifier Bt1491, recently designated BtuJ1) showing cartoon representation of the β-jellyroll barrel with B_12_ in magenta lines (PDB code 9FCT), and right is the corresponding schematic β-sheet topology diagram. The inset depicts the genomic neighbourhood of Bt1491, showing its proximity to genes encoding *btuB, btuC, btuD*, and other predicted vitamin B_12_ transport proteins. D) Pairwise sequence identity matrix for the DUF4465 proteins shown in panel A selected for assessment of B_12_-binding function.

### Determination of B_12_-binding to DUF4465 Family Proteins

To investigate the ubiquity of B_12_ binding within the DUF4465 protein family, we cloned genes representing the signal peptide-truncated forms of five representative proteins (D5EK51, Pan, Fluta, 656, and Cylst) and expressed them in *E. coli*. All five proteins bound B_12_ during purification, as indicated by their persistent pink colour when bound to the metal affinity column, and upon elution from a size-exclusion column, each protein retained an intense pink colour and exhibited the characteristic shift to slightly smaller hydrodynamic radius when complexed with B_12_ compared with the apo form (Supplementary Figure 1).

### Determination of the structure of the D5EK51-B_12_ Complex

D5EK51-B_12_ crystals belonged to the space group P3_2_21, with three D5EK51-B_12_ complexes in the asymmetric unit. The structure revealing the detailed D5EK51-B_12_ interactions was solved by molecular replacement and refined to 1.85□Å resolution, yielding R-work and R-free values of 0.19 and 0.212, respectively, for a model with high stereochemical quality. Crystallographic and refinement statistics are summarized in Supplementary Table 1.

### Structural Features and B_12_ Binding Mechanism

The D5EK51 protein adopts an augmented β-jellyroll fold. This fold consists of eight β-strands arranged into two antiparallel four-stranded β-sheets, BIDG and CHEF, as shown in Figure 2A, B. In D5EK51, the β-jellyroll architecture is extended by five additional antiparallel β-strands at the N-terminus (WXYZA) and an additional α-helix inserted within loop DE. This structural arrangement is consistent with other recently solved DUF4465 family members, BtuJ1 and BtuJ2 (PDB codes: 9FCT and 9I2L)^24^.

**Figure 2.**
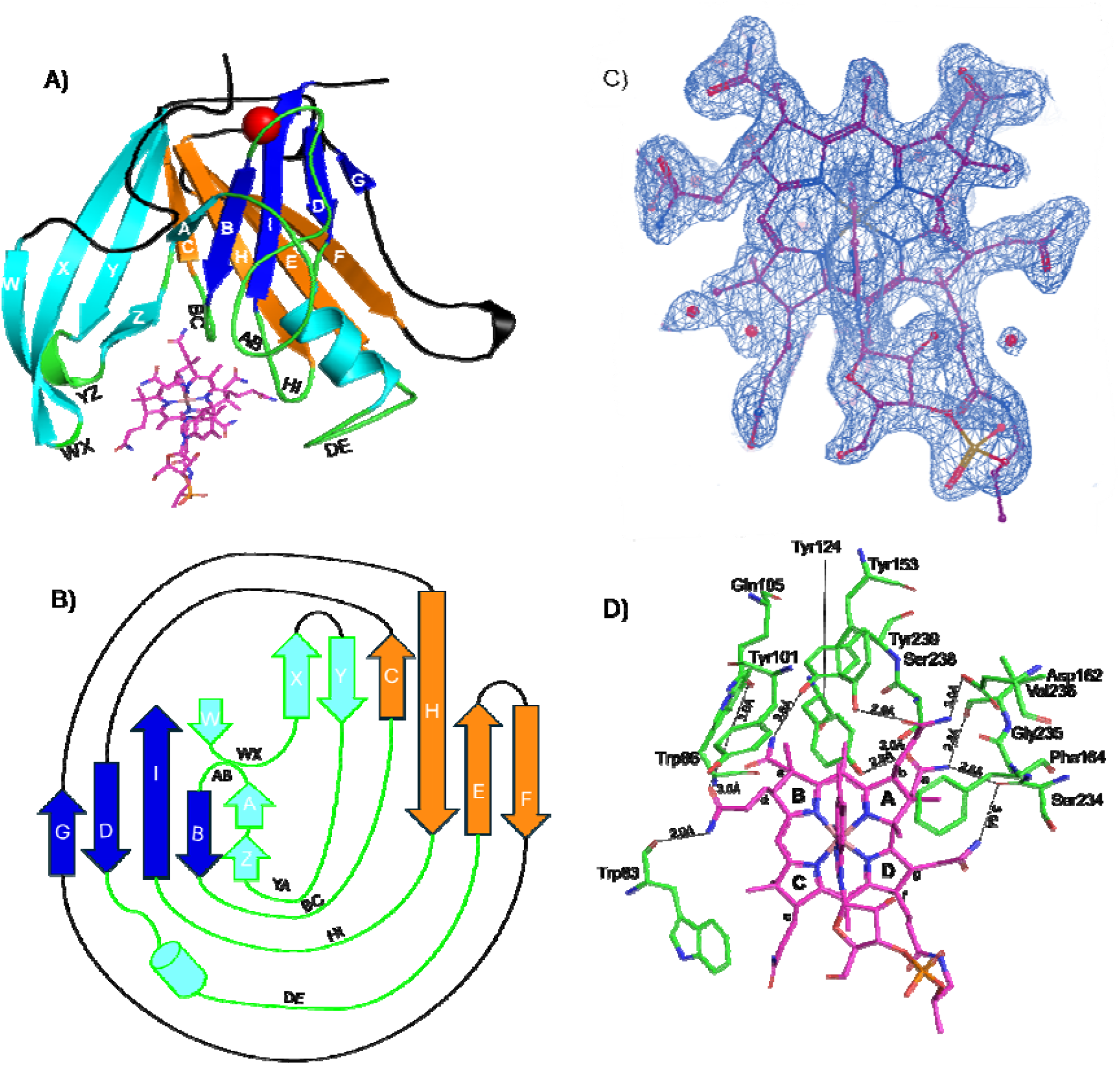
Structural features of the D5EK51–B_12_ complex. A) Overall cartoon representation of the D5EK51 structure, highlighting the augmented β-jellyroll fold characteristic of DUF4465 family proteins. The two β-sheets are labelled according to convention: BIDG (blue) and CHEF (orange). Five additional β-strands (WXYZA) at the N-terminus and a loop-associated α-helix (loop DE) are also present. Loops are names according to the β-strands they link together; hence loop HI connects β-strands H and I. Loops forming the B_12_-binding site are shown in green. Cyanocobalamin is depicted in magenta, and a stabilizing calcium ion is shown as a red sphere. B) Topology diagram of the D5EK51 secondary structure. The β-jellyroll fold consists of two antiparallel four-stranded β-sheets with additional N-terminal β-strands (WXYZA) and an α-helix (loop DE), consistent with other DUF4465 family proteins. C) 2Fo−Fc electron density map (blue mesh) of the B_12_ ligand contoured at 1.0σ, showing well-resolved density for the corrin ring and side chains. Ordered water molecules are also visible and contribute to ligand binding. D) Close-up view of the B_12_-binding pocket showing the hydrogen bonding network (dashed lines). Eleven hydrogen bonds are formed between D5EK51 and B_12_, involving both main-chain and side-chain atoms. These interactions predominantly occur near the A-ring of the corrin ring, stabilizing B_12_ within a hydrophobic cavity composed of aromatic residues shown in green.

Cyanocobalamin is well-resolved within each molecule of the asymmetric unit (Figure 2C). The binding pocket is formed by six loops conserved among DUF4465 proteins and is primarily lined with aromatic hydrophobic residues, including tyrosine, phenylalanine and tryptophan. These aromatic residues not only accommodate the corrin ring but also form contacts with its side chains, facilitating tight binding (Figure 2D). In addition, a network of eleven hydrogen bonds is established between D5EK51 and the B_12_ side chains, involving both backbone and side-chain atoms.

D5EK51 shares 28% and 21% sequence identity with BtuJ1 and BtuJ2 and their structures align with root-mean-square deviation (RMSD) of Å 3.548 and 1.518 Å, respectively. The HI loop and the DE loop/helix (named after the β-strands connected) are highly conserved among the three structures, both in their conformation and in the identity of B_12_-coordinating residues (Supplementary Figure 2). These loops contain the conserved B_12_-binding tyrosine’s, on loop HI (Y245 in BtuJ1 numbering) and within the α-helix of loop DE (Y146), highlighting a preserved functional core of the DUF4465 family.

Surface conservation mapping further supports this conclusion (Figure 3A). Key aromatic residues within the binding pocket are conserved across homologs, particularly those in loops YZ, AB, BC, HI, and DE. The HI loop (GTPAYF) is especially conserved, with all residues present at greater than 70% frequency and the critical B_12_-binding tyrosine retained in 97% of homologs. Sequence motif alignment was used to create a sequence logo (Figure 3B) and extension of this analysis representatives of all eight clade (Figure 3C) strongly supports B_12_ binding as a conserved function across the DUF4465 family.

**Figure 3.**
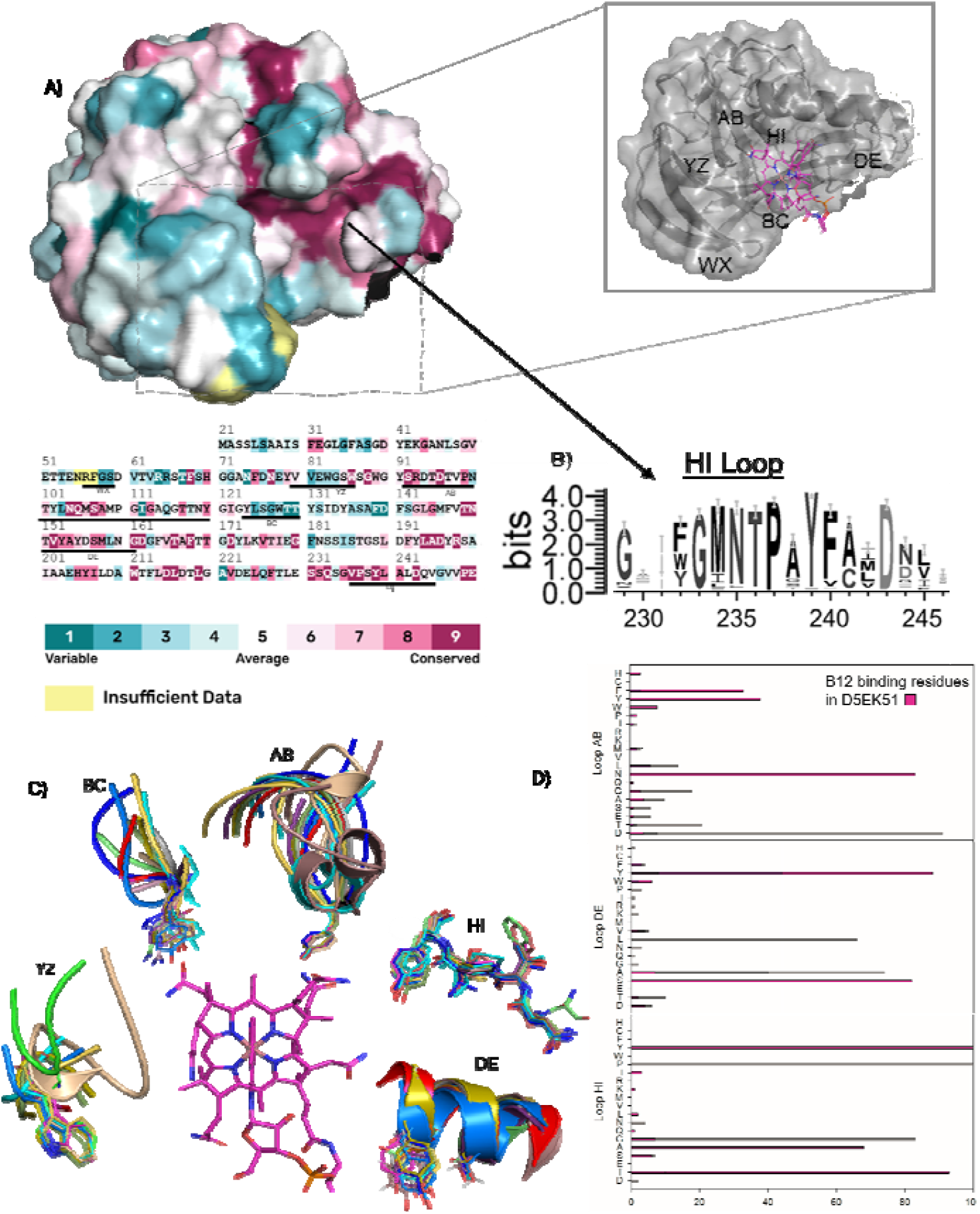
ConSurf analysis of D5EK51 reveals evolutionary conservation of the B_12_-binding pocket. A) Surface representation of the experimentally determined structure colored by residue conservation (teal indicates variable and pink conserved residues), highlighting the conserved B_12_-binding pocket (boxed region). Below is a multiple sequence alignment of representative homologs with conservation scores (1–9) mapped to the structure. B) Sequence logo for residues predicted on HI loop, showing a conserved NTPAY motif. C) Structural alignment of AlphaFold models of DUF4465 homologs from different bacterial clades bound to cobalamin (magenta), revealing a conserved ligand-binding orientation. D) Residue frequency diagram for the three B_12_-binding loops.

### Genomic analysis of Genetic Context of DUF4465 encoding genes

We identified two major genomic clusters of DUF4465 proteins. The first, comprising approximately 900 members, showed frequent co-localization with transport-associated domains: 20% co-located with ABC-type transporters (PF00005), 40% with TonB-dependent β-barrels (PF00593), 55% with their associated plug domains (PF07715), and 40% with β-propeller domain proteins (PF16819). The second cluster, containing 195 DUF4465 members, exhibited even stronger associations, with 80–95% co-occurring with these transport systems and periplasmic binding proteins. In total, 84 DUF4465 proteins lacked neighboring Pfam-annotations, suggesting that these values may underestimate the true frequency of such associations. Together, these patterns indicate that DUF4465 proteins are frequently genomically linked to vitamin B_12_ transport systems, supporting their role in B_12_ acquisition.

## Materials and Methods

### Molecular biology

Fragments corresponding to the soluble domains of the DUF4465 proteins, lacking the predicted lipoprotein motif, were inserted into a modified pET3a vector resulting in constructs encoding N-terminal hexahistidine tagged proteins produced via the T7 promotor. The plasmids were validated by sequencing. The seven stared proteins in Figure 1A were produced in this way.

### Protein expression

*E. coli* BL21(DE3) cells were used for protein production. Transformants were selected using 100 μg/ml ampicillin. Single colonies were used to inoculate 50 mL Luria broth (100 μg/ml ampicillin) and cultured overnight at 37°C with shaking. 10 mL of overnight culture was used to inoculate 800 mL fresh media and cultivated until the culture reached mid-log phase, measured with an OD_600_ value of 0.6. At this point the culture was cooled and induced with 0.5 mM Isopropyl β-d-1-thiogalactopyranoside (IPTG). The induced culture was incubated at 20°C overnight, with shaking. Bacteria were collected by centrifugation, and the obtained pellets kept at –20°C until required.

### Protein purification

The bacterial pellet from 400 mL culture was resuspended in 100 mL of binding buffer (20 mM Tris-Cl, 150 mM NaCl, 30 mM imidazole, and 1 mM tris(2-carboxyethyl) phosphine (TCEP), and sonicated on ice. The obtained cell lysates were clarified by ultracentrifugation at 100,000xg for 40 minutes at 4°C. The resulting supernatant was applied to a HisTrap HP column containing 5 mL of chelating Sepharose. The His-tagged DUF4465 protein was then eluted in 5 mL fractions with elution buffer (20 mM Tris-Cl, 150 mM NaCl, 500 mM imidazole, and 1 mM TCEP). Subsequently, the protein mixture was separated using size exclusion chromatography (SEC) on an S200 HiLoad column equilibrated with SEC buffer (20mM Tris-Cl, 150mM NaCl, 1mM TCEP). The eluted protein fractions were then concentrated and applied to a size-exclusion chromatography column (Superdex200, 16/16) in buffer 20 mM Tris-HCl, pH 7.5, 150 mM NaCl, 1 mM TCEP. The fractions corresponding to the expected elution volumes for the purified DUF domains were collected, flash-frozen in liquid nitrogen, and stored at -80 °C. To produce DUF proteins in complex with B_12_ excess cyanocobalamin was added to the cell lysate before the affinity column. Unbound B_12_ was removed during the HisTrap HP column washing step.

### Protein crystallization and structure determination

Protein was concentrated to 20 mg/ml for crystallization trials using sitting drop vapour diffusion trials set up a Mosquito Crystallization robot (TTP Labtech) using the commercial screens JCSG+, Structure 1 and 2, Morpheus, MembranePlus, Proplex, LMB, MIDAS+ (Molecular Dimensions and Hampton Research). Crystallization plates were incubated at 20°C. Crystals of D5EK51 grew in MIDAS+ and Memgold2 commercial screens. Crystallization was achieved using the MIDAS+ condition 34-1, comprising 35% poly(acrylic acid sodium salt) 2100, 0.1□M HEPES (pH 7.5), and 0.2□M ammonium sulfate. Crystals were mounted and washed in cryoprotectant augmented with 20% glycerol and then flash frozen in liquid nitrogen in loops for data collection. Diffraction data were collected using the DECTRIS EIGER X 9M at ESRF beamline BM30A. Structure determination, refinement and validation used molecular replacement with an AlphaFold^27,28^-generated model of D5EK51 within the CCP4 suite^29^. All molecular graphics were prepared using PyMOL (The PyMOL Molecular Graphics System, Version 2.0 Schrödinger, LLC)^30^.

### Sequence similarity network and genome neighbourhood analysis

Multiple sequence alignments were obtained using MUSCLE5 and MAAFT^31–33^. Proteins belonging to the IPR027828 family (DUF4465) were analysed using the Enzyme Function Initiative Enzyme Similarity Tool (EFI-EST) and Genome Neighbourhood Tool (EFI-GNT) (https://efi.igb.illinois.edu). An SSN was generated by submitting the InterPro family identifier IPR027828, retrieving UniProtKB sequences clustered at 90% sequence identity to reduce redundancy. An alignment score threshold of 15 (corresponding to c. 40% sequence identity) was used to define edges, yielding two major clusters. For genomic context analysis, representative sequences from each connected component were submitted to EFI-GNT, examining genes within ±10 open reading frames (ORFs). Gene neighbourhoods were annotated based on EFI-generated diagrams and UniProt functional descriptions. Analysis of the signal sequences were performed by signalP 6.0^26^.

## Acknowledgements

This work was supported by the Biotechnology and Biological Sciences Research Council (BBSRC) under Grant BB/X001946/1. We are grateful for beamtime at ESRF (Grenoble) beamline BM30A. We acknowledge the European Synchrotron Radiation Facility (ESRF) for provision of synchrotron radiation facilities under proposal number mx2606/mx2698 and we would like to thank beamline staff for assistance and support in using beamline BM30A. The authors would also like to thank Diamond Light Source for beamtime (proposal mx38316), and the staff of beamlines I02, I03 and I24 for assistance with crystal testing and data collection.

## Data availability

The structure and structure factors for DUF4465 protein D5EK51 is deposited in the protein databank with code 9HMU.

## References

1. Sokolovskaya, O. M., Shelton, A. N. & Taga, M. E. Sharing vitamins: Cobamides unveil microbial interactions. Science (1979) 369, (2020).

2. Bassford, P. J., Bradbeer, C., Kadner, R. J. & Schnaitman, C. A. Transport of vitamin B12 in tonB mutants of Escherichia coli. J Bacteriol 128, 242–7 (1976).

3. Shelton, A. N. et al. Uneven distribution of cobamide biosynthesis and dependence in bacteria predicted by comparative genomics. ISME J 13, 789–804 (2019).

4. Degnan, P. H., Barry, N. A., Mok, K. C., Taga, M. E. & Goodman, A. L. Human Gut Microbes Use Multiple Transporters to Distinguish Vitamin B12 Analogs and Compete in the Gut. Cell Host Microbe 15, 47–57 (2014).

5. Magnúsdóttir, S., Ravcheev, D., de Crécy-Lagard, V. & Thiele, I. Systematic genome assessment of B-vitamin biosynthesis suggests co-operation among gut microbes. Front Genet 6, (2015).

6. Croft, M. T., Lawrence, A. D., Raux-Deery, E., Warren, M. J. & Smith, A. G. Algae acquire vitamin B12 through a symbiotic relationship with bacteria. Nature 438, 90–93 (2005).

7. Degnan, P. H., Taga, M. E. & Goodman, A. L. Vitamin B12 as a Modulator of Gut Microbial Ecology. Cell Metab 20, 769–778 (2014).

8. Kräutler, B. Biochemistry of B12-Cofactors in Human Metabolism. in 323–346 (2012). doi:10.1007/978-94-007-2199-9_17.

9. Roth, J., Lawrence, J. & Bobik, T. COBALAMIN (COENZYME B12): Synthesis and Biological Significance. Annu Rev Microbiol 50, 137–181 (1996).

10. Smith, A. D., Warren, M. J. & Refsum, H. Vitamin B12. in 215–279 (2018). doi:10.1016/bs.afnr.2017.11.005.

11. Rodionov, D. A., Vitreschak, A. G., Mironov, A. A. & Gelfand, M. S. Comparative Genomics of the Vitamin B12 Metabolism and Regulation in Prokaryotes. Journal of Biological Chemistry 278, 41148–41159 (2003).

12. Zhang, Y., Rodionov, D. A., Gelfand, M. S. & Gladyshev, V. N. Comparative genomic analyses of nickel, cobalt and vitamin B12 utilization. BMC Genomics 10, 78 (2009).

13. Villa, E. A. & Escalante-Semerena, J. C. Corrinoid salvaging and cobamide remodeling in bacteria and archaea. J Bacteriol 206, (2024).

14. Helliwell, K. E. et al. Cyanobacteria and Eukaryotic Algae Use Different Chemical Variants of Vitamin B12. Current Biology 26, 999–1008 (2016).

15. Magnúsdóttir, S., Ravcheev, D., de Crécy-Lagard, V. & Thiele, I. Systematic genome assessment of B-vitamin biosynthesis suggests co-operation among gut microbes. Front Genet 6, (2015).

16. Magnúsdóttir, S., Ravcheev, D., de Crécy-Lagard, V. & Thiele, I. Systematic genome assessment of B-vitamin biosynthesis suggests co-operation among gut microbes. Front Genet 6, (2015).

17. Helliwell, K. E. The roles of B vitamins in phytoplankton nutrition: new perspectives and prospects. New Phytologist 216, 62–68 (2017).

18. Shultis, D. D., Purdy, M. D., Banchs, C. N. & Wiener, M. C. Outer Membrane Active Transport: Structure of the BtuB:TonB Complex. Science (1979) 312, 1396–1399 (2006). x19.

19. Korkhov, V. M., Mireku, S. A., Veprintsev, D. B. & Locher, K. P. Structure of AMP-PNP–bound BtuCD and mechanism of ATP-powered vitamin B12 transport by BtuCD–F. Nat Struct Mol Biol 21, 1097–1099 (2014).

20. Cadieux, N. et al. Identification of the Periplasmic Cobalamin-Binding Protein BtuF of Escherichia coli. J Bacteriol 184, 706–717 (2002).

21. Wexler, A. G. et al. Human gut Bacteroides capture vitamin B12 via cell surface-exposed lipoproteins. Elife 7, (2018).

22. Putnam, E. E. et al. Gut Commensal Bacteroidetes Encode a Novel Class of Vitamin B12-Binding Proteins. mBio 13, (2022).

23. Abellon-Ruiz, J. et al. BtuB TonB-dependent transporters and BtuG surface lipoproteins form stable complexes for vitamin B12 uptake in gut Bacteroides. Nat Commun 14, 4714 (2023).

24. Juodeikis, R. et al. Extracellular Vesicle-Linked Vitamin B12 Acquisition via Novel Binding Proteins in Bacteroides Thetaiotaomicron. (2025) doi:10.1101/2025.07.31.667871.25.

25. Abellon-Ruiz, J. et al. BtuJ1, a Novel Surface-Exposed B12 -Binding Protein in Bacteroidetes , Functions as an Extracellular Vitamin Reservoir That Enhances Fitness. (2025) doi:10.1101/2025.07.01.662609.

26. Almagro Armenteros, J. J. et al. SignalP 5.0 improves signal peptide predictions using deep neural networks. Nat Biotechnol 37, 420–423 (2019).

27. Varadi, M. et al. AlphaFold Protein Structure Database in 2024: providing structure coverage for over 214 million protein sequences. Nucleic Acids Res 52, D368–D375 (2024).

28. Jumper, J. et al. Highly accurate protein structure prediction with AlphaFold. Nature 596, 583–589 (2021).

29. Agirre, J. et al. The CCP4 suite: integrative software for macromolecular crystallography. Acta Crystallogr D Struct Biol 79, 449–461 (2023).

30. Schrödinger, L. The PyMOL Molecular Graphics System, Version 2.0. Preprint at (2025).

31. Kuraku, S., Zmasek, C. M., Nishimura, O. & Katoh, K. aLeaves facilitates on-demand exploration of metazoan gene family trees on MAFFT sequence alignment server with enhanced interactivity. Nucleic Acids Res 41, W22–W28 (2013).

32. Katoh, K., Rozewicki, J. & Yamada, K. D. MAFFT online service: multiple sequence alignment, interactive sequence choice and visualization. Brief Bioinform 20, 1160–1166 (2019).

33. Edgar, R. C. High-accuracy alignment ensembles enable unbiased assessments of sequence homology and phylogeny. Preprint at 10.1101/2021.06.20.449169 (2021).

