## Supplementary figures and images for "Evolutionarily divergent DUF4465 domains have a common vitamin B_12_-binding function"

### Supplemental Table 1

**Supplementary Table 1.** Crystallographic and refinement statistics for D5EK51

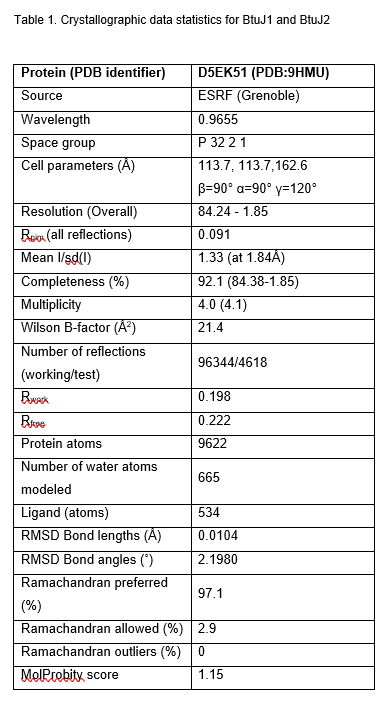
