## Supplemental Figure 1 for "Evolutionarily divergent DUF4465 domains have a common vitamin B_12_-binding function"

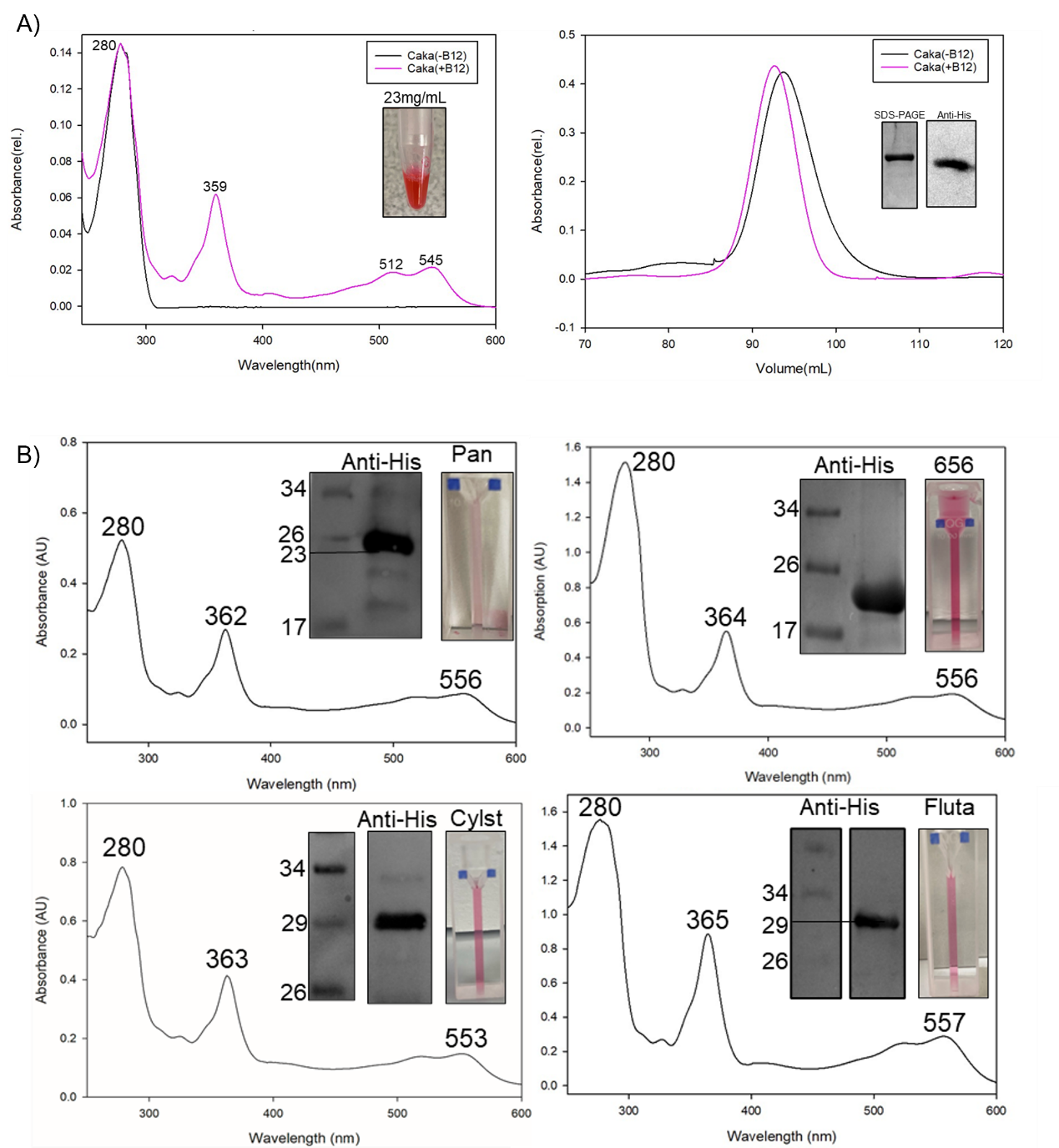

**Supplementary Figure 1.** Spectroscopic and chromatographic characterization of B_12_-binding to DUF4465 proteins. A) UV-visible absorbance spectra of purified D5EK51 alone (black) and in complex with cyanocobalamin (pink). The D5EK51–B_12_ (Caca-B_12_) complex exhibits distinct absorbance features at 359 nm, and 545 nm, consistent with B_12_-binding, while the unbound protein shows only the protein 280 nm absorbance. The inset shows the deep pink coloration of the D5EK51–B_12_ complex at 23 mg/mL, indicative of bound cobalt-containing cobalamin. Size-exclusion chromatograms of D5EK51 purified with (pink) or without (black) B_12_ on a Superdex 200 HiLoad column. Both samples elute at c. 95 mL, corresponding to the monomeric form of D5EK51 (26.9 kDa). The B_12_-complex elutes slightly earleir than the protein alone as seen previously for BtuJ1 and BtuJ2. SDS-PAGE and anti-His Western blot (inset) gel confirms the purity of D5EK51. B) Purification and spectral analysis of each DUF4465 protein showing characteristic B_12_ absorption peaks (uv–vis spectra) and anti-His immunoblot confirmation of protein expression. Insets show purified protein-B_12_ complex (Pan, 656, Cylst and Fluta proteins).
