## Supplemental Figure 2 for "Evolutionarily divergent DUF4465 domains have a common vitamin B_12_-binding function"

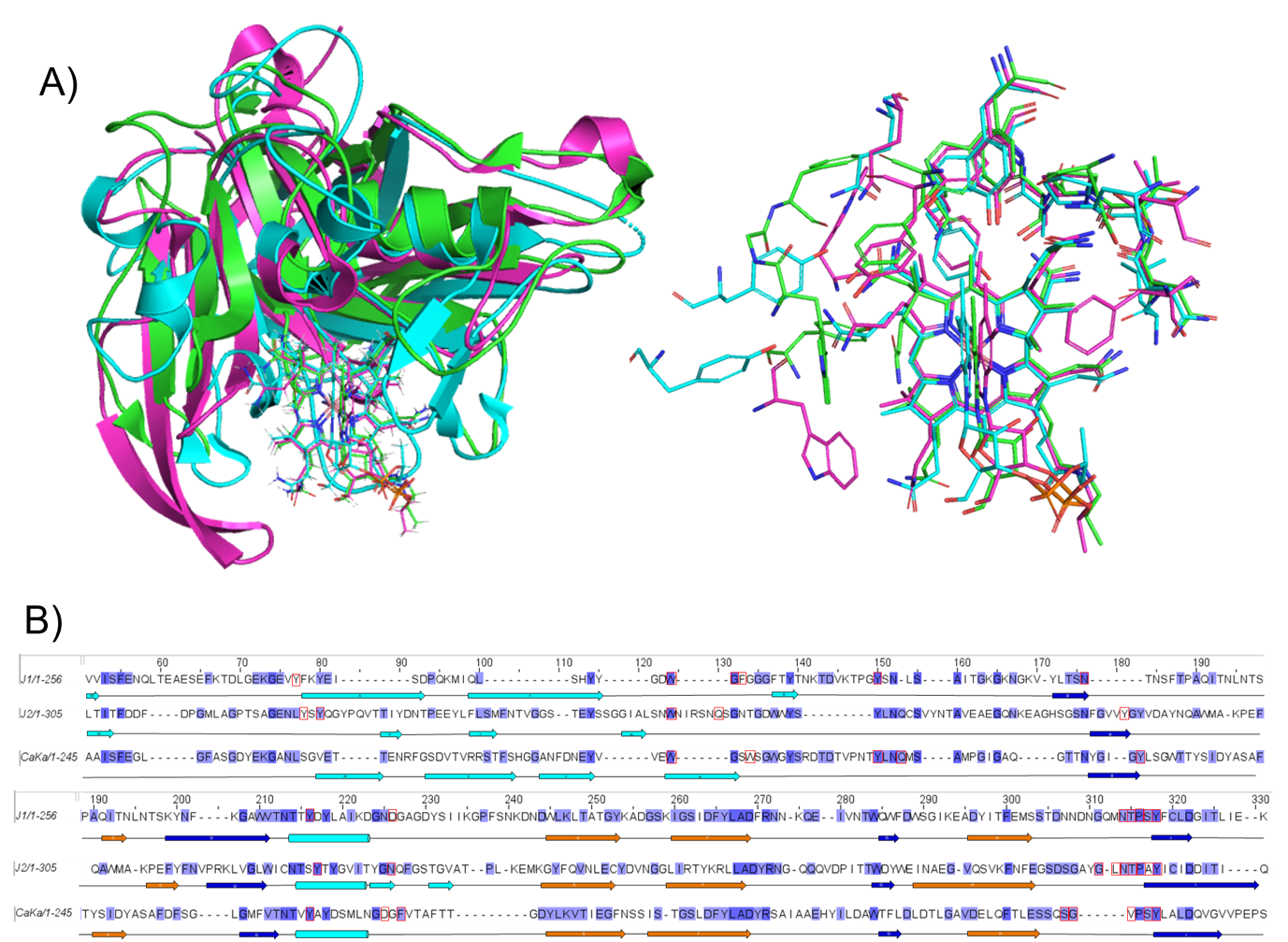

**Supplementary Figure 2.** Structural comparison and sequence alignment of D5EK51, BtuJ1 and BtuJ2. A) Structural superposition of three DUF4465 proteins in complex with B_12_—BtuJ1 (green), BtuJ2 (cyan), and D5EK51 (magenta). All three structures adopt the conserved β-jellyroll fold and coordinate B_12_ within a similarly positioned pocket. Top right is a close-up view of the B_12_-binding site which reveals conservation of aromatic and hydrogen bonding residues responsible for B_12_-coordination. Side chains from all three structures show near-identical orientation around the A ring of B_12_ molecule, highlighting a shared ligand recognition strategy. B) Structure-based sequence alignment of BtuJ1, BtuJ2, and D5EK51, with secondary structure elements annotated. Conserved residues involved in B_12_ binding are indicated inside of red boxes, and secondary structure elements are shown as arrows (β-strands) and cylinders (α-helices).
